# Reconstitution of minimal motility system based on *Spiroplasma* swimming by expressing two bacterial actins in synthetic minimal bacterium

**DOI:** 10.1101/2021.11.16.468548

**Authors:** Hana Kiyama, Shigeyuki Kakizawa, Yuya Sasajima, Yuhei O Tahara, Makoto Miyata

## Abstract

Motility is one of the most important features of life, but its evolutionary origin remains unknown. In this study, we focused on *Spiroplasma*, commensal, or parasitic bacteria. They swim by switching the helicity of a ribbon-like cytoskeleton that comprises six proteins, each of which evolved from a nucleosidase and bacterial actin called MreB. We expressed these proteins in a synthetic, non-motile minimal bacterium, JCVI-syn3.0B, whose reduced genome was computer-designed and chemically synthesized. The synthetic bacterium exhibited swimming motility with features characteristic of *Spiroplasma* swimming. Moreover, some combinations of the two proteins produced a helical cell shape and swimming, suggesting that the swimming originated from the differentiation and coupling of bacterial actins, and we also obtained a minimal system for motility of the synthetic bacterium.

**One-Sentence Summary:** The minimal system comprised two bacterial actins that provided cell helicity and swimming to the synthetic minimal bacterium.

---

Motility is observed in various phyla and is arguably one of the major determinants of survival. If we focus on the force-generating units of cell motility, all cell motilities reported to date can be classified into 18 mechanisms (*1*). Generally, the direct evolutionary ancestor of the individual mechanisms cannot be identified, probably because several of these have been in existence for a long time. However, it is possible to discuss their origin and evolution. Cell motility is considered to originate from the rather small movements of housekeeping proteins which are amplified and transmitted to the cell outside, possibly because of the accumulation of mutations. However, this process has not yet been experimentally demonstrated. Class Mollicutes are parasitic or commensal bacteria that are characterized by a small genome (*2, 3*). Interestingly, there are three unique motility mechanisms in Mollicutes (*4–6*). It is likely that when the phylum Firmicutes evolved to stop peptidoglycan synthesis, they also stopped flagellar motility, which depends on the peptidoglycan layer, and then acquired unique motility (*1, 5*). In one of the three types of motilities, when *Spiroplasma* swims, they thrust water by switching the handedness of their helicity (*4, 7–9*). These schemes are completely different from those of the spirochete, a group of bacteria with helical cells. The helical shape of *Spiroplasma* is likely determined by a ribbon-like cytoskeleton, which comprises fibril protein evolved from nucleosidases (*10–12*), and five classes of *Spiroplasma* MreBs evolved from MreB, the bacterial actin (*12–15*). Here, we refer to *Spiroplasma* MreBs as SMreBs because they are distantly related to MreBs found in walled-bacteria (*13, 16, 17*). The helicity of the ribbon is determined by the fibril protein, but the mechanism of helicity switching is unknown.

The synthetic bacterium JCVI-syn3.0 was established by J. Craig Venter Institute (JCVI) in 2016 as a combination of a cell of *Mycoplasma capricolum* and a genome designed based on *Mycoplasma mycoides*. Both *Mycoplasma* species belong to *the Spiroplasma* clade, one of four Mollicutes clades (*2*). It has a fast growth rate, which is beneficial for genome manipulation, roughly spherical morphology, and no motility (*18, 19*). JCVI-syn3.0B (syn3B) has 19 genes returned from *M. mycoides* for better growth (*18*). In this study, we reconstituted *Spiroplasma* swimming in syn3B by adding seven genes, and identified the minimal gene set for *Spiroplasma* cell helicity and swimming.

## Results

### Reconstitution of *Spiroplasma* swimming in syn3B

We focused on *Spiroplasma eriocheiris*, an actively swimming pathogen in crustaceans (*14*). Seven genes that are likely related to swimming are encoded in four loci in the genome: fibril, five classes of SMreB, and a non-annotated conserved gene (*4, 13, 16, 17*). We assembled these genes into an 8.4 kb DNA fragment, and incorporated it into the syn3B genome using the *Cre/loxP* system (Figs. 1A and Fig. S1, Table S1) (*20, 21*). An active promoter in syn3B, Ptuf, was inserted upstream of the gene cluster. Surprisingly, under optical microscopy, 48% of the syn3B cells exhibited a filamentous cell shape and active movements, presumably accompanied by force generation, and 13% had a helical shape and swimming motility (Fig. 1B, Movie S1). We named this construct syn3Bsw. The width and pitch of the cell helices analyzed by optical and electron microscopy were slightly different from those of *Spiroplasma* cells (Fig. 1C). If we focus on cells that are partially bound to the glass, we can observe that a free part of the cell was rotating with some reversals (Fig. 1D, Movie S2), meaning that helicity switching causes helix rotation in syn3Bsw, similar to *Spiroplasma* swimming. Next, we analyzed the helices and handedness of cell images in each frame of the swimming video (Fig. 1E, F). The handedness of the cell helix differed depending on its axial position, and the helicity changed over time. Further, we measured the movement and rotation speed of the helix from the part where it appeared to move along the cell axis smoothly. The helix movement and rotation speeds were 8.2 ± 3.7 μm/s and 11.6 ± 4.8 /s (n=10) for syn3sw, not significantly different from 8.8 ± 2.8 μm/s and 12.0 ± 3.6 /s (n=16) for *Spiroplasma*. In the cryo-electron microscopy (EM) image of syn3Bsw cells, filaments running along the axis were observed in the inner part of the curvature, similar to *Spiroplasma* cells (Fig. S2). The filaments recovered from syn3Bsw cells exhibited chained ring structures characteristic of the fibril filament from *Spiroplasma* cells (Figs 1G and S3) (*10, 11, 14*). The periodicities were 8.5±0.9 nm (n=41) and 8.7±1.0 nm (n=48) for *Spiroplasma* and syn3Bsw without significant differences (*p* = 0.45 by Student’s *t-test*). In addition, electrophoretic and mass spectrometric analyses of cell lysates indicated that fibrils and all SMreBs were expressed in syn3Bsw cells (Fig. S4, Table S2). These results indicate that the expression of *Spiroplasma* proteins inside syn3Bsw cells resulted in the formation of internal filaments that reconstituted helical shape, helicity switching, and swimming.

**Fig. 1.**
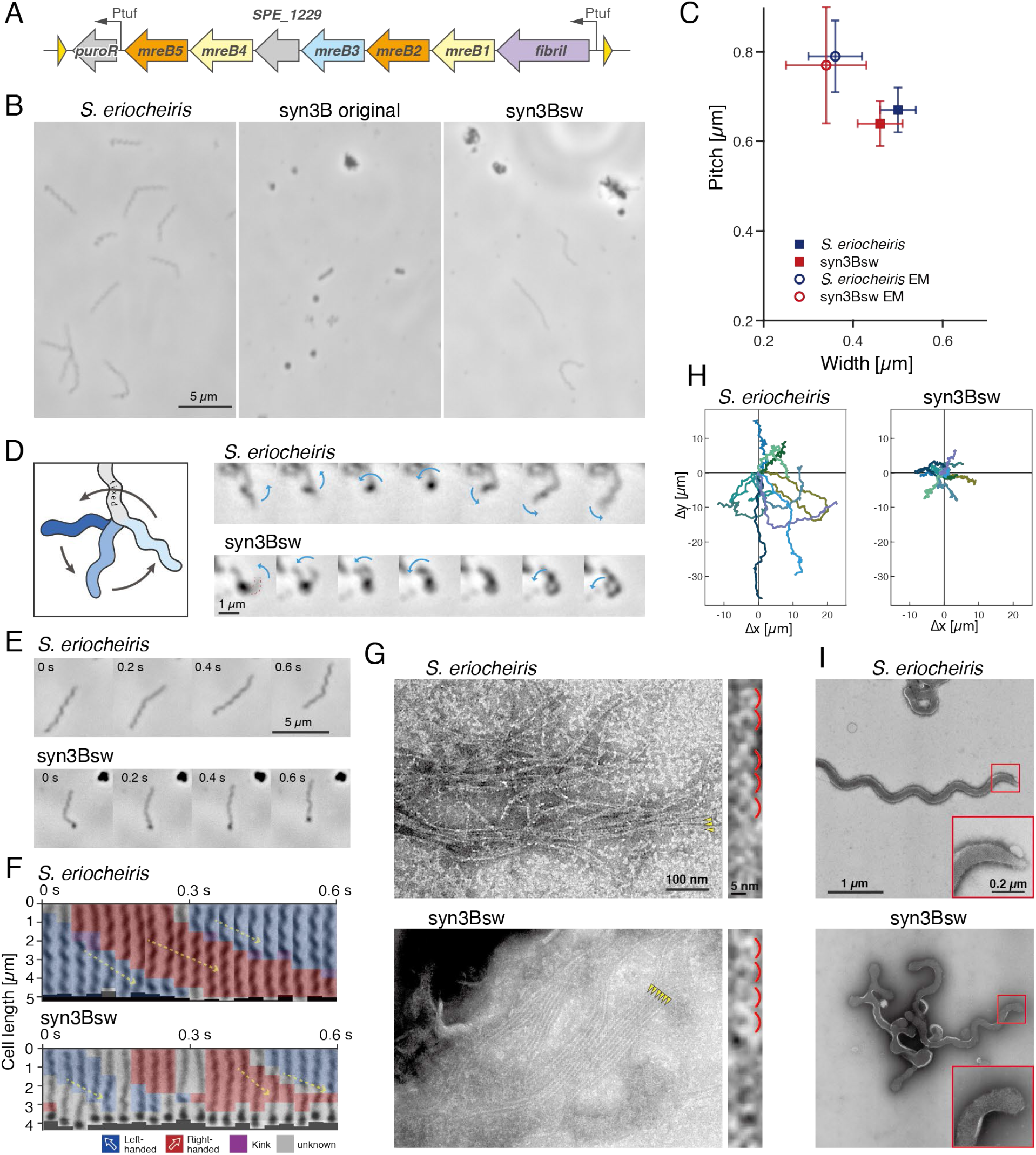
Reconstitution of Spiroplasma swimming in syn3B by expressing seven genes. (**A**) A DNA fragment transferred between two *loxP* sites of syn3B, including seven genes from *Spiroplasma* and a puromycin resistance gene, “*puroR*.” A non-annotated gene, *SPE_1229* is indicated by a gray arrow. Ptuf and *loxP* sites are denoted by black and yellow triangles, respectively. (**B**) Field cell images of three strains are indicated on the top. In syn3Bsw, DNA fragment illustrated in (A) is inserted into the genome by Cre/*loxP* system. The cells were observed by phase contrast microscopy. (**C**) Distribution of cell helicity parameters measured by optical and electron microscopy. The *p* values by Student’s *t*-test between Spiroplasma and syn3Bsw were 0.008, 0.002, 0.59, and 0.28 for pitch and width in optical microscopy, and EM, respectively. (**D**) Rotational behaviors of freely moving parts of *Spiroplasma* and syn3Bsw cells. A schematic is illustrated in the left. The cell is fixed to the glass through the light gray part, and the blue part rotates. Consecutive video frames are shown for every 0.03 s. A rotational behavior of the free part is marked by blue arrows. The rotating part in syn3Bsw is marked by a red broken line. (**E**) Consecutive video frames of swimming cells for every 0.2 s. (**F**) Change in helicity analyzed for videos shown in (E). The cell images were straightened and analyzed by Image J, and then colored for their handedness. Smooth traveling helix is marked by a yellow arrow. (**G**) Negative-staining EM images of filaments recovered from *Spiroplasma* and syn3Bsw cells. Filaments are marked by yellow triangles. Magnified images are illustrated in the right with marks for ring structures, characteristic for fibril filament. (**H**) Traces of a pole of ten cells for 10 s colored differently. (**I**) Cell images under negative-staining EM images of *Spiroplasma* and syn3Bsw cells. A cell pole is magnified as inset.

### Differences in swimming between syn3Bsw and *Spiroplasma*

The speeds of helix movement and rotation were not significantly different between syn3Bsw and *Spiroplasma* (Fig. 1F). However, the trajectory of the cells over 10 s indicated that syn3Bsw could not travel long distances, unlike *Spiroplasma* (Fig. 1H). The reason can be seen in the time course of helicity switching, indicating little continuity in the rotation that hampers long-distance traveling (Fig. 1F). This may be caused by a lack of cooperativity in the helicity switching that generates helix rotation. EM images of syn3Bsw cells did not show the tapered pole, including an inner architecture called “dumbbell,” unlike *Spiroplasma* cells (Fig. 1I) (*14*), suggesting that the tapered pole made by unknown proteins plays a role in continuous helicity switching of the ribbon.

### Role of Component Proteins

To examine the role of each protein, we produced and analyzed constructs in which each protein was not expressed (Figs. 2 A and B, Movie S3). To avoid affecting gene expression by the alteration in the DNA and RNA structures, we introduced nonsense mutations to the 8th–22nd codons of each structural gene (Fig. S1). We confirmed by electrophoresis that the target proteins were no longer expressed in the mutant cells (Fig. S5). No significant differences from syn3Bsw were observed in cell structures and behaviors for five of the six constructs (Fig. 2A). However, in the construct missing SMreB5, the helix width was 0.64 ± 0.13 μm, significantly larger than that of syn3Bsw in half of the filamentous cells, and the cells moved but did not swim. The distinctive features of the lack of SMreB5 are consistent with a previous observation that *Spiroplasma citri* lost helicity and swimming because of the absence of SMreB5 (*13*). These results suggest that the seven proteins have some redundant roles in helix formation and swimming.

**Fig. 2.**
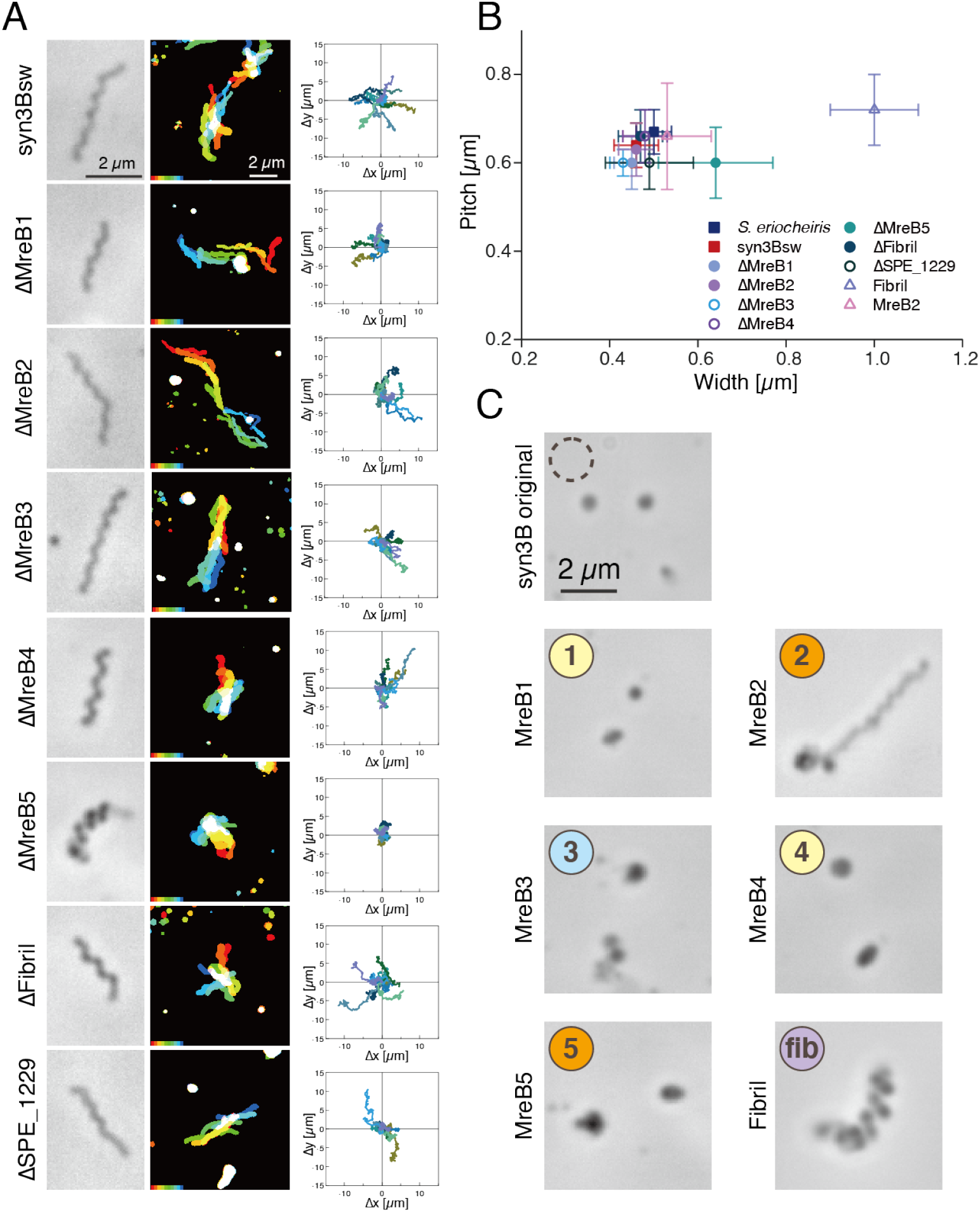
Role of individual proteins in syn3 swimming. (**A**) Structure and behaviors of cells lacking one of seven proteins from syn3sw. For each construct, phase-contrast cell image (left), integrated cell images every 1 s for 10 s with colors changing from red to blue (middle), and traces of a pole of ten cells for 10 s (right) are indicated. (B) Distribution of cell helicity parameters for individual constructs analyzed with optical microscopy. (C) Phase-contrast image of cells expressing single *Spiroplasma* protein marked by “fib” and number of SMreBs. The original syn3B is marked by a broken circle.

We then examined syn3B constructs expressing each protein (Fig. 2C, Movie S4). The cells expressing only fibril protein formed a helical cell shape with a pitch of 0.72 ± 0.08 μm and a width of 1.0 ± 0.10 μm, which is wider than *Spiroplasma* cells. The pitch of the helix is in good agreement with the number of isolated fibrils, which is consistent with the fact that fibrils are a major component of the ribbon (*10–12, 14*). The cells expressing SMreB2 formed filamentous morphology, and some of them formed helices with a variety of pitches of 0.66 ± 0.12μm. The cells expressing only SMreBs1, 3, 4, or 5 did not show differences in cell shape from the original syn3B.

### Expressing a pair of SMreBs

Further, we analyzed the shapes and behaviors of cells expressing ten combinations of SMreB protein pairs (Fig. 3A, Fig. S1). As five classes of SMreB can be divided into three groups based on their amino acid sequence: 5–2, 4–1, and 3 (*13, 16, 17*), we will discuss the results based on this classification. In the pairs of SMreBs selected from the 5–2 and 4–1 groups, surprisingly, the cells of the 5–1 and 5–4 combinations exhibited helix formation and movements, and some cells showed swimming like syn3Bsw, with occurrence frequencies comparable to syn3Bsw (Figs. 3, B and C, Movie S5). Cells of 2–1 showed a filamentous morphology and movements. The cells of the 2–4 combination showed filamentous morphology, but were immotile; however, a few in several hundred cells showed movement. In the combinations of one of the 5–2 or 4–1 groups paired with 3, cells of 3–2 formed a right-handed helix (Fig. 3D, Movie S6). In the combinations of 3–1, 3–4, and 3–5, the cells did not show differences from the original syn3B. In the combinations of the same group, 5–2 and 4–1 cells were filamentous, and 4–1 cells rarely formed a short, right-handed helix.

**Fig. 3.**
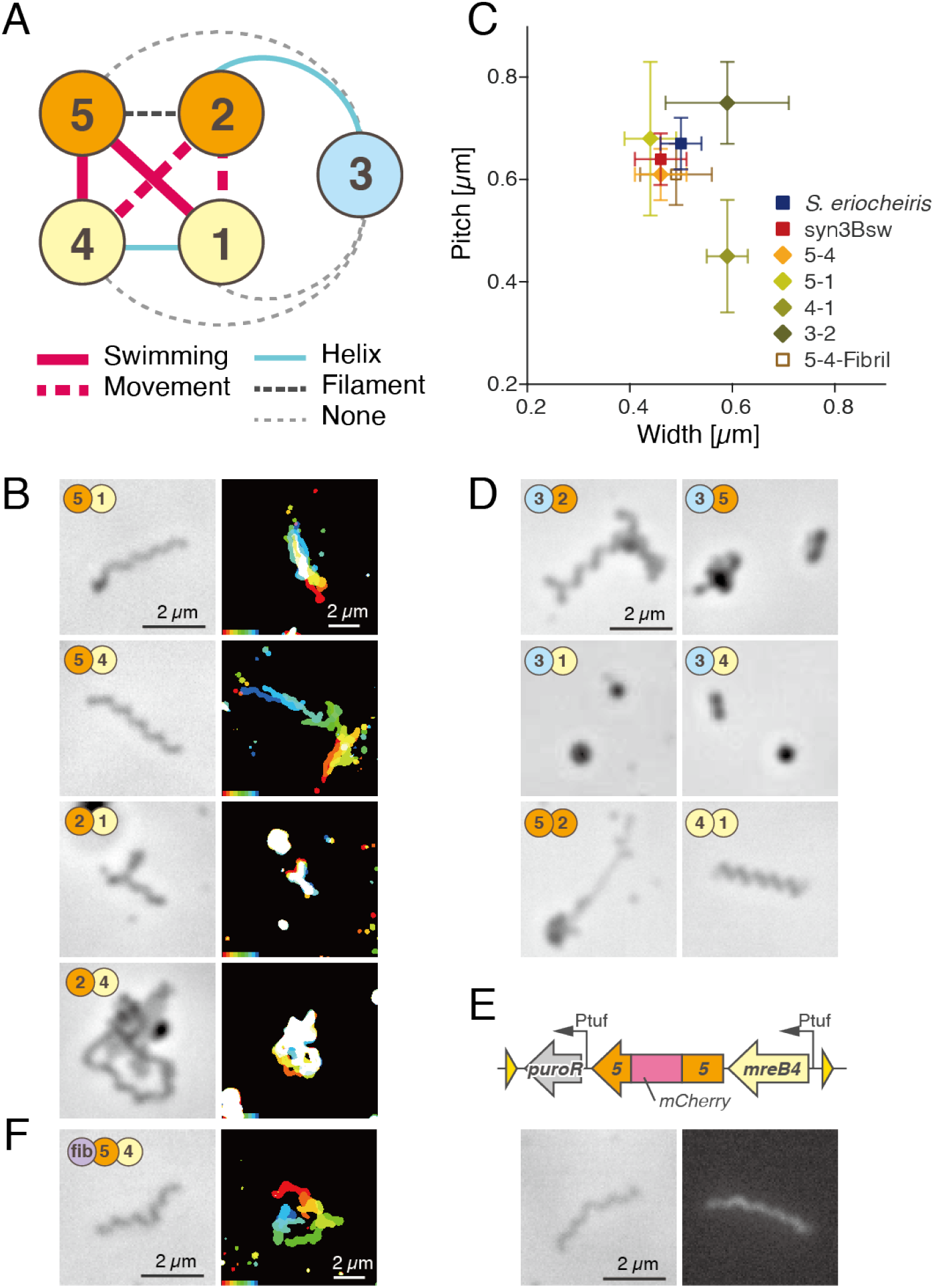
Morphology and behaviors of syn3B cells expressing pair of SMreB proteins. (**A**) Schematic of SMreB combinations with protein groups. Each SMreB is presented by a numbered circle with a group color. The characters that result in syn3B cells by gene expression are presented by line formats. (**B**) Image (left) and behaviors (right) of syn3B cells expressing pair of SMreBs. Cells of four constructs presented here showed movements. (**C**) Distribution of parameters for cell helicity. (**D**) Phase-contrast image of cells expressing other combinations of protein pairs. Six pairs did not show movements. (**E**) SMreB5 localization in cell expressing SMreBs 4 and 5. Schematic of integrated genes is illustrated (upper). mCherry gene is inserted into the C-terminal side of tyrosine residue at the 218th position. Phase-contrast and fluorescence images are illustrated (lower). (**F**) Image (left) and behaviors (right) of syn3B cells expressing SMreBs 4, 5, and fibril.

In the construct of 5–4, we fused the fluorescent protein mCherry into SMreB 5 and 4 at a position suggested by previous studies (Figs. 3E and Fig. S1, Movie S7) (*22*). The cells expressing SMreB5 fused with mCherry showed a helical cell shape and swimming, as observed in the 5–4 cells. Fluorescence was observed throughout the cell, suggesting that SMreB5 filaments were formed along the entire cell axis. In addition, this result indicated that mCherry fusion did not interfere with the functions of SMreB5. The 5–4 cells with mCherry fusion to SMreB4 did not exhibit conspicuous helicity. Even helical cells found in hundreds of cells did not show any movement. To clarify the roles of fibril, a major component of the ribbon structure, we analyzed cells expressing fibrils in addition to SMreBs 4 and 5 (Fig. 3F, Movie S8). The differences between the presence and absence of fibril protein were subtle in the analyses conducted in this study.

## Discussion

MreB belongs to the actin superfamily and forms a short antiparallel double-strand filament based on ATP energy (*23, 24*). It has the ability to sense the curvature of the peripheral structures and serves to guide the bacterial peptidoglycan synthase to positions required for the synthesis (*25*). Isolated SMreBs also form fibers similar to those of MreB (*13, 26*). Our results indicate that helix formation and force generation of *Spiroplasma* occur by the interaction between different SMreBs. The mechanism can be explained as follows (Fig. 4): Protofilaments made of proteins belonging to either the SMreB 5–2 or 4–1 groups are aligned along the cell axis and bound together. If the unit length in each protofilament is different, some curvature is induced in the double strand, resulting in helix formation. If these protofilaments undergo a local length change at different times using ATP energy, the curvature changes like a bi-metallic strip, resulting in helicity switching (*4*). The length change may be related to polymerization and depolymerization in terms of the change in axial distance between the subunits. Remarkably, the differences in the amino acid sequence between SMreBs 5–2 and 4–1 groups in *S. eriocheiris* range from 29.1–31.6% if similar amino acid pairs are excluded (*16*). These small differences suggest that the ancestors of SMreB may have acquired stability, helicity, and switching after the accidental acquisition of different properties. Specifically, it may represent the moment when a slight structural change in a housekeeping protein is amplified by an accidental accumulation of mutations, leading to motility. The reason for the existence of as many as five SMreBs, even though two proteins are capable of acquiring helicity and force generation, is unclear. It may be advantageous for efficient and robust swimming, possibly in different environments, or for chemotaxis. The participation of fibrils can be explained in a similar manner. To the best of our knowledge, the motility system comprising only two actin superfamily proteins is the smallest system established till date (*1*). Therefore, we may call this a “minimal motile cell.”

**Fig. 4.**
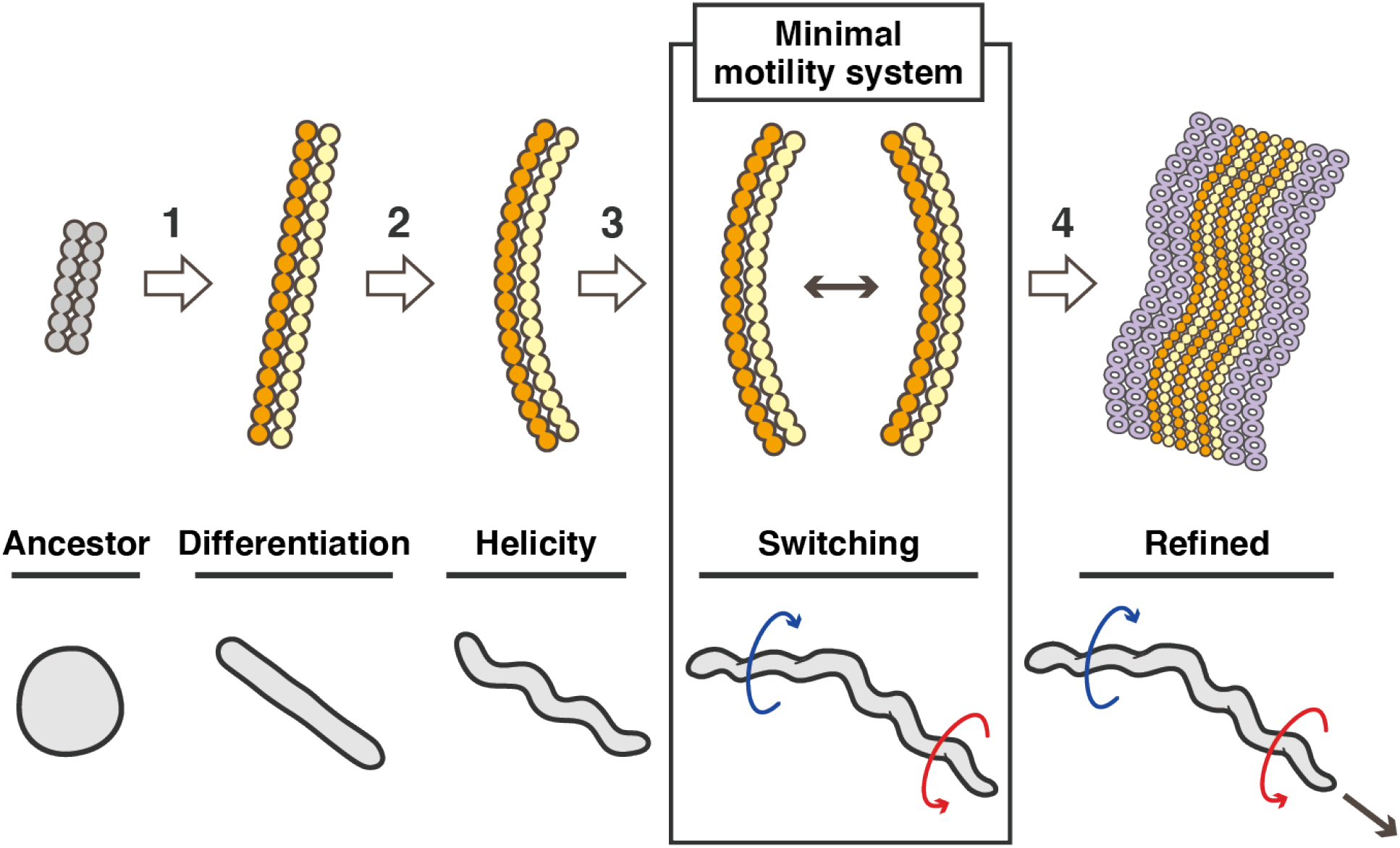
Development and mechanism of *Spiroplasma* swimming. The swimming mechanism may be acquired through four steps as denoted by white arrows. Step 1: The MreB protein derived from walled bacteria differentiated into two classes with different characters by accumulated mutations. Association of heterogeneous protofilaments allowed stable filament formation. Step 2: Small differences of protofilaments in length generated curvature, resulting in helicity of the heterogenous filament. Step 3: Change in protofilament length caused by ATP energy induces change in curvature, causing helicity switching. Then, the initial stage, in other words, the minimal motility system was acquired. Step 4: The acquired swimming was refined to be equipped by five classes of SMreBs, fibril, dumbbell structure, etc. Corresponding cell morphology and behaviors are presented (lower).

Here, we used JCVI-syn3B as the experimental platform (*18, 19*). Because the genes of synthetic bacteria are derived from organisms related to *Spiroplasma*, it remains possible that factors derived from this near-minimal synthetic bacterium, such as proteins related to cell division (*18*) or the composition and physical properties of the cell membrane, are essential for helix formation and swimming. Thus, completely controlling factors linked to cell functions is still a future challenge. Nevertheless, the results of this study demonstrate that syn3B is a good system for studying cell functions and their evolution.

## Supporting information

Plasmid sequences

## Acknowledgments

We acknowledge Prof. John Glass, Prof. Kim Wise, and JCVI for their helpful inputs on JCVI-syn3.0B and improving the manuscript, Ikuko Fujiwara, Takuma Toyonaga for helpful discussions, and Junko Shiomi and Tomomi Shimonaka for technical assistance.

## Author contributions

Conceptualization: HK, SK, MM; Methodology: SK, YS, Investigation: HK, YS, Visualization: HK, YS, YOT; Writing – original draft: HK, MM; Writing – review and editing: HK, SK, MM; Funding acquisition: MM, HK. Competing interests: The authors declare no conflicts of interest.

## Data and materials availability

All data are available in the main text or supplementary materials.

