## Supplemental materials for "Reconstitution of minimal motility system based on *Spiroplasma* swimming by expressing two bacterial actins in synthetic minimal bacterium"

**This PDF file includes:**

Materials and Methods
Figs. S1 to S5
Tables S1 to S2
Captions for Movies S1 to S8
Captions for Data S1

**Other Supplementary Materials for this manuscript include the following:**

Movies S1 to S8
Data S1

### Materials and Methods

**Bacterial strains and culture conditions.** JCVI-syn3.0B (GenBank: CP069345.1), *Spiroplasma eriocheiris* (TDA-040725-5T), *and Escherichia coli* (DH5α) for DNA manipulation were cultured in SP4 (*18, 19*), SP4, and LB media at 37 °C, 30 °C, and 37 °C, respectively.

**Plasmid construction.** The *Spiroplasma* genome was isolated as previously described (*27*). The plasmid utilized to transform JCVI-syn3.0B to obtain syn3Bsw (pSeW001) was constructed as follows (Fig. S1). Focused *Spiroplasma* DNA regions, *puroR* gene, and vector fragment were amplified from the *Spiroplasma* genome DNA and pSD079 DNA (*21*) as five PCR products, using the primer sets listed in Table S1. The DNA fragments were assembled using the In-Fusion® HD Cloning Kit (Takara Bio Inc. Kusatsu, Japan). pSeW002 was constructed by replacing the upstream region of the first gene, *a fibril* in pSeW001 with a Ptuf fragment (promoter from the EF-Tu gene) amplified from pSD079. pSeW102, pSeW202, pSeW302, pSeW402, pSeW502, pSeW602, and pSeW702 were modified to introduce nonsense mutations in individual genes. The plasmids utilized to construct other strains were modified from pSeW005, which was constructed by the process described above, using pSeW002 as the PCR template. Ptuf or Pspi (Spiralin promoter from *Spiroplasma*) (*21*) were inserted at the 5' end of the first open reading frame (ORF). All the DNA fragments were verified for DNA sequences.

**Transformation and cell preparation.** The transformation of JCVI-syn3.0B was performed as previously described (*28*) with two modifications: (i) the entire process was scaled down to 1/15 and (ii) the mixture of cells and DNA was kept on ice for 10 min to increase its transformation efficiency. The transformant colonies were picked and inoculated into a 200 µl SP4 medium containing 3 µg/ml puromycin, cultured for 18–24 h, and confirmed for transformation by PCR. The cells were cultured in a liquid medium a few more times, with an inoculation of 100–500 dilution, and then frozen as stock. To analyze the cells, the frozen stock was inoculated at 100–500 dilution in SP4 medium with puromycin, and grown for 20–24 h at 37 °C. Cultures at an optical density of 0.03 at 620 nm were used for the analyses of JCVI-syn3.0B and *S. eriocheiris*.

**Optical microscopy and protein profiling.** The cultured cells of syn3B and *Spiroplasma* were analyzed in a 0.5 × SP4 medium diluted with PBS, containing 0.5% methylcellulose and 0.5 mg/mL BSA. If necessary, the cell density was adjusted by centrifugation at 11, 000 × g at 10 °C for 10 min, followed by suspension with the diluted medium. The cell suspension was inserted into a tunnel slide (*14, 27, 29, 30*), and observed using an inverted microscope IX71 (Olympus, Tokyo, Japan) equipped with a UPlanSApo 100 × 1.4 numerical aperture (NA) Ph3 and complementary metal-oxide-semiconductor (CMOS) camera, DMK33UX174 (The Imaging Source Asia Co., Ltd. Taipei, Taiwan). The videos were analyzed by ImageJ ver.1.53f51 (Fiji) using plugins, MTrackJ, empirical gradient threshold (EGT), and a color footprinting macro (*31*). Profiling and identification of proteins in cells was performed as previously described (*14, 32, 33*).

**Electron microscopy.** To observe the intact cells, cultured cells were collected by centrifugation, suspended to a 10-fold density of the original in the medium, and fixed using 0.5% glutaraldehyde for 5 min at 25 °C. After quenching with 500 mM Tris-HCl pH7.5, the cells were collected by centrifugation, washed, and suspended in PBS to a 40-fold density of the original.

The cell suspension was placed on a carbon-coated grid for 5 min, removed, rinsed with PBS thrice, and then stained with 2% phosphotungstic acid for 60 s. To observe the internal structure, the cell suspension was treated with PBS containing 1% Triton X-100, 0.1 mg/mL DNase, 1 mM $MgCl_2$, and 1mM PMSF (phenylmethylsulfonyl fluoride) for 10 min at 4 °C, and centrifuged at 20,000 × g for 30 min at 4 °C. The pellet was suspended in PBS to a 160-fold density of the original, placed on the EM grid for 2 min, and stained with 2% phosphotungstic acid for 60 s. Images were acquired using a JEM1010 EM (JEOL, Akishima, Japan) equipped with a FastScan-F214(T) charge-coupled device (CCD) camera (TVIPS, Gauting, Germany). For cryoEM, the cultured cells were collected and suspended to a 10-fold density of the original, and frozen as described previously (*34*). The images were captured using a Talos F200C EM (Thermo Fisher Scientific, Waltham, MA, USA) equipped with a 4k × 4k Ceta CMOS camera (Thermo). The images were analyzed using ImageJ software.

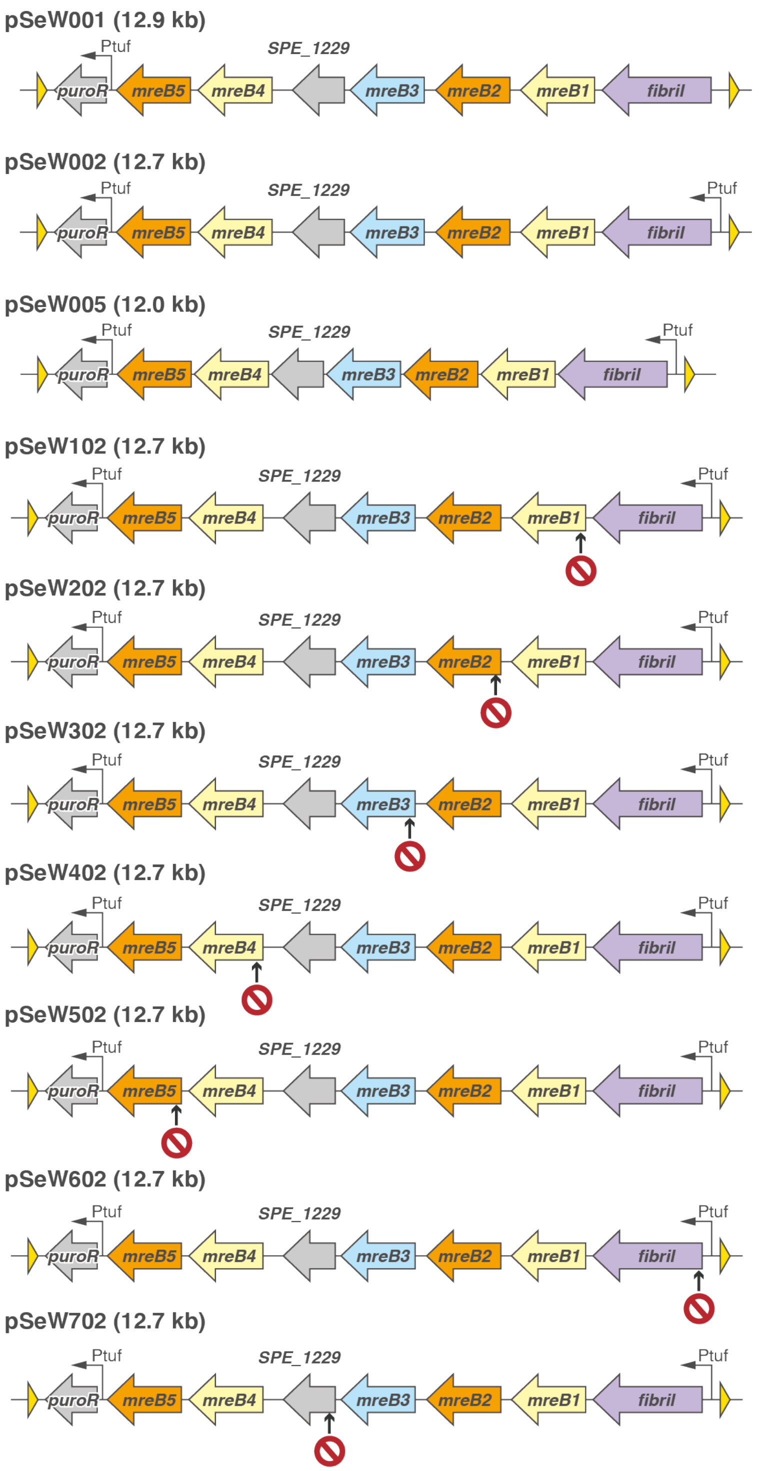
pSeW001 (12.9 kb)
Ptuf
SPE_1229
puroR
mreB5
mreB4
mreB3
mreB2
mreB1
fibril
pSeW002 (12.7 kb)
pSeW005 (12.0 kb)
pSeW102 (12.7 kb)
pSeW202 (12.7 kb)
pSeW302 (12.7 kb)
pSeW402 (12.7 kb)
pSeW502 (12.7 kb)
pSeW602 (12.7 kb)
pSeW702 (12.7 kb)

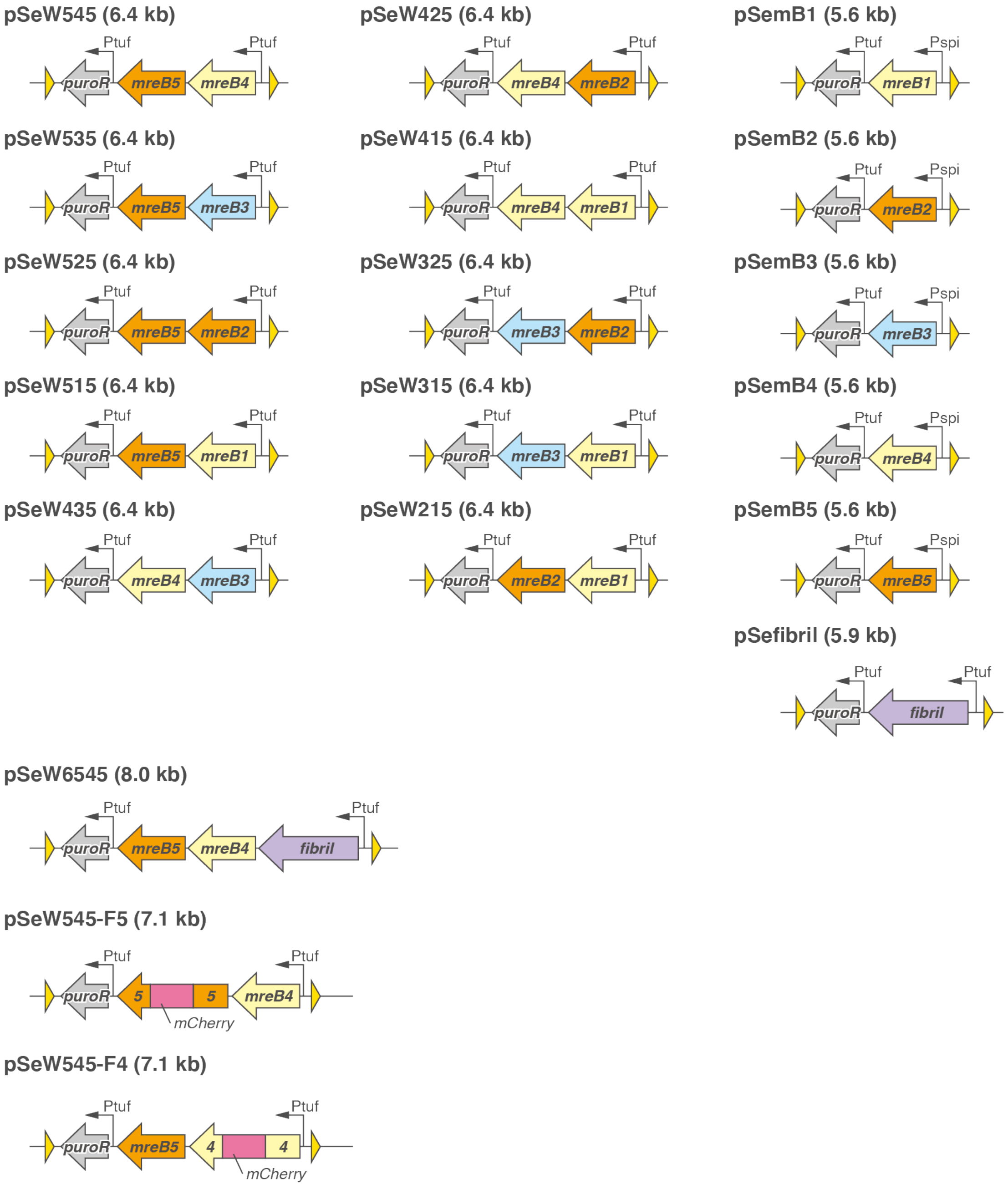


**Fig. S1.**

Schematic presentation for DNA constructs used in this study.

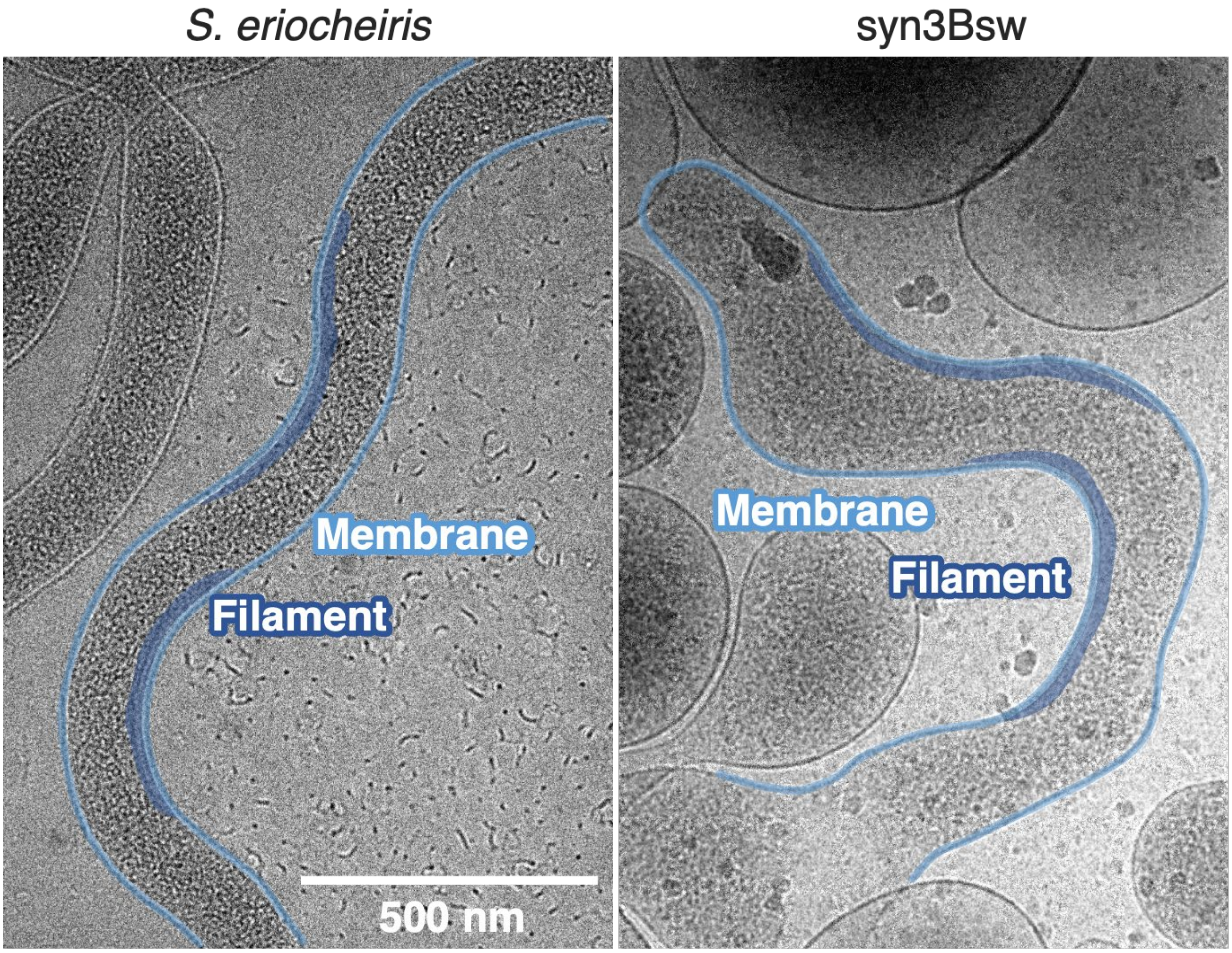


**Fig. S2.**
Cell images under cryo electron microscopy. The cell membrane and filamentous structures are colored light and dark blue, respectively.

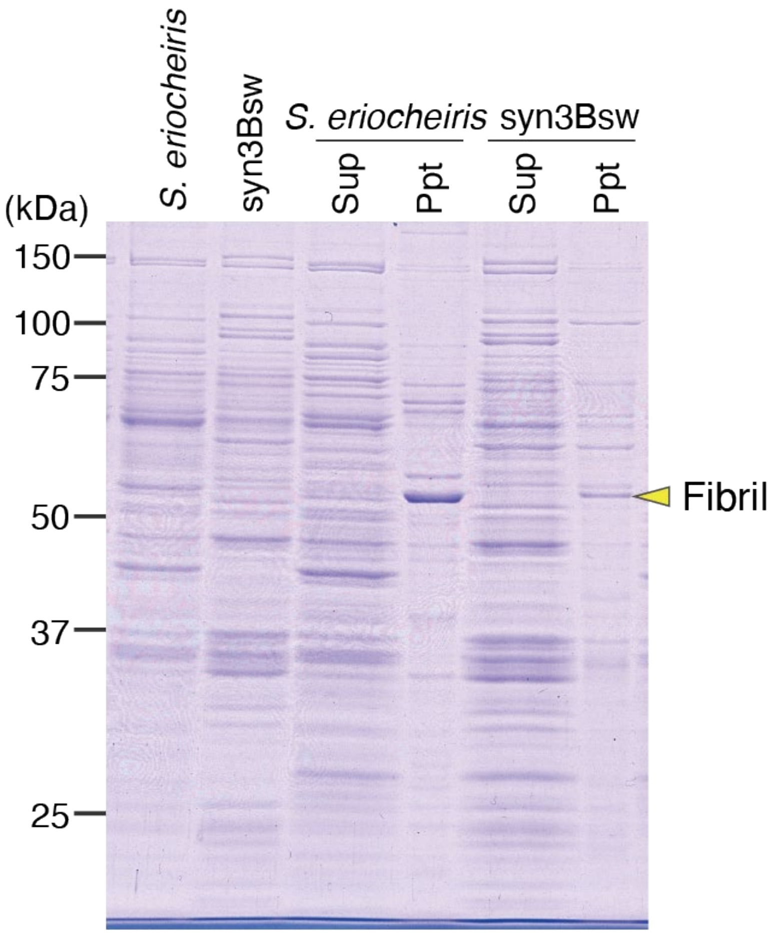


**Fig. S3.**

Identification of fibril protein in syn3Bsw. The cultured cells were washed, lyzed , fractionated by centrifugation, and analyzed by SDS-10% PAGE. The entire cell lysates are illustrated in the left two lanes. Fibril protein is marked. Protein bands were stained by Coomassie Brilliant Blue.

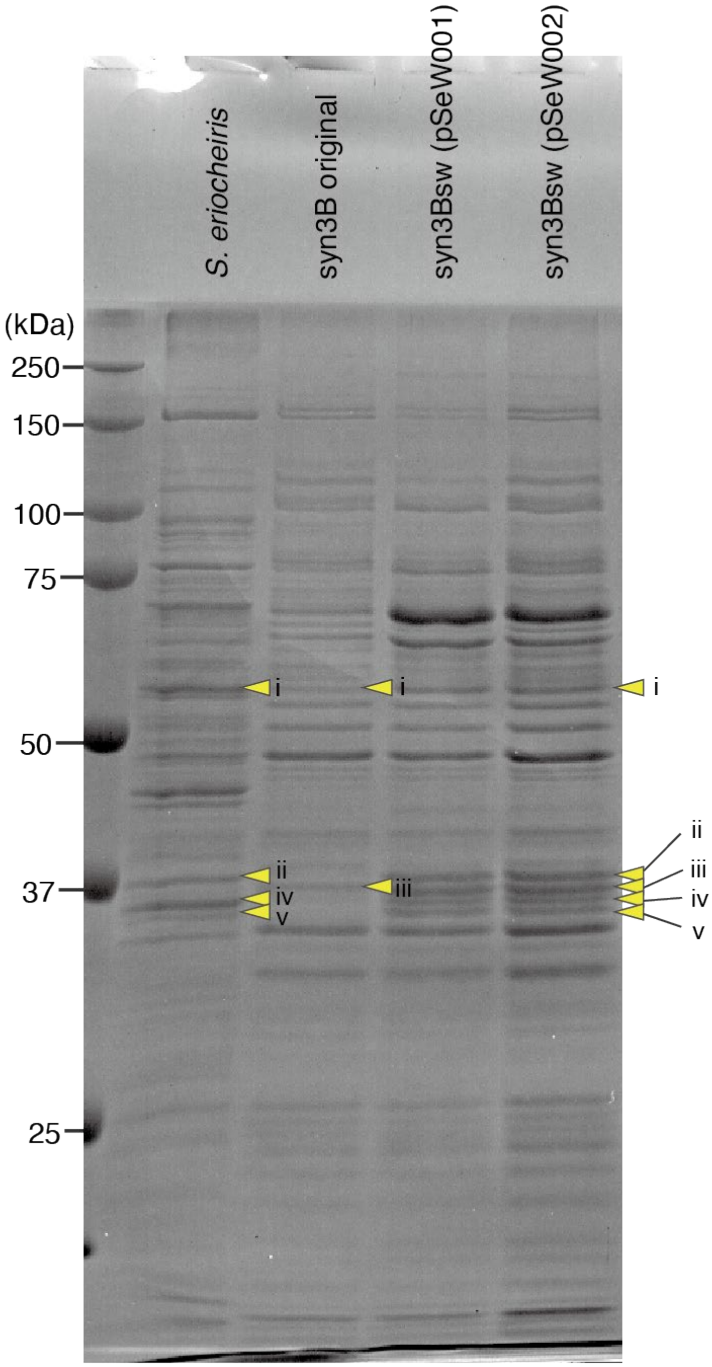


**Figure S4.**

Whole protein profiles of cell lysates visualized by SDS-12.5% PAGE. Constructs are indicated on the top, and focused protein bands are marked by a yellow triangle with a number. Protein bands were stained by Coomassie Brilliant Blue.

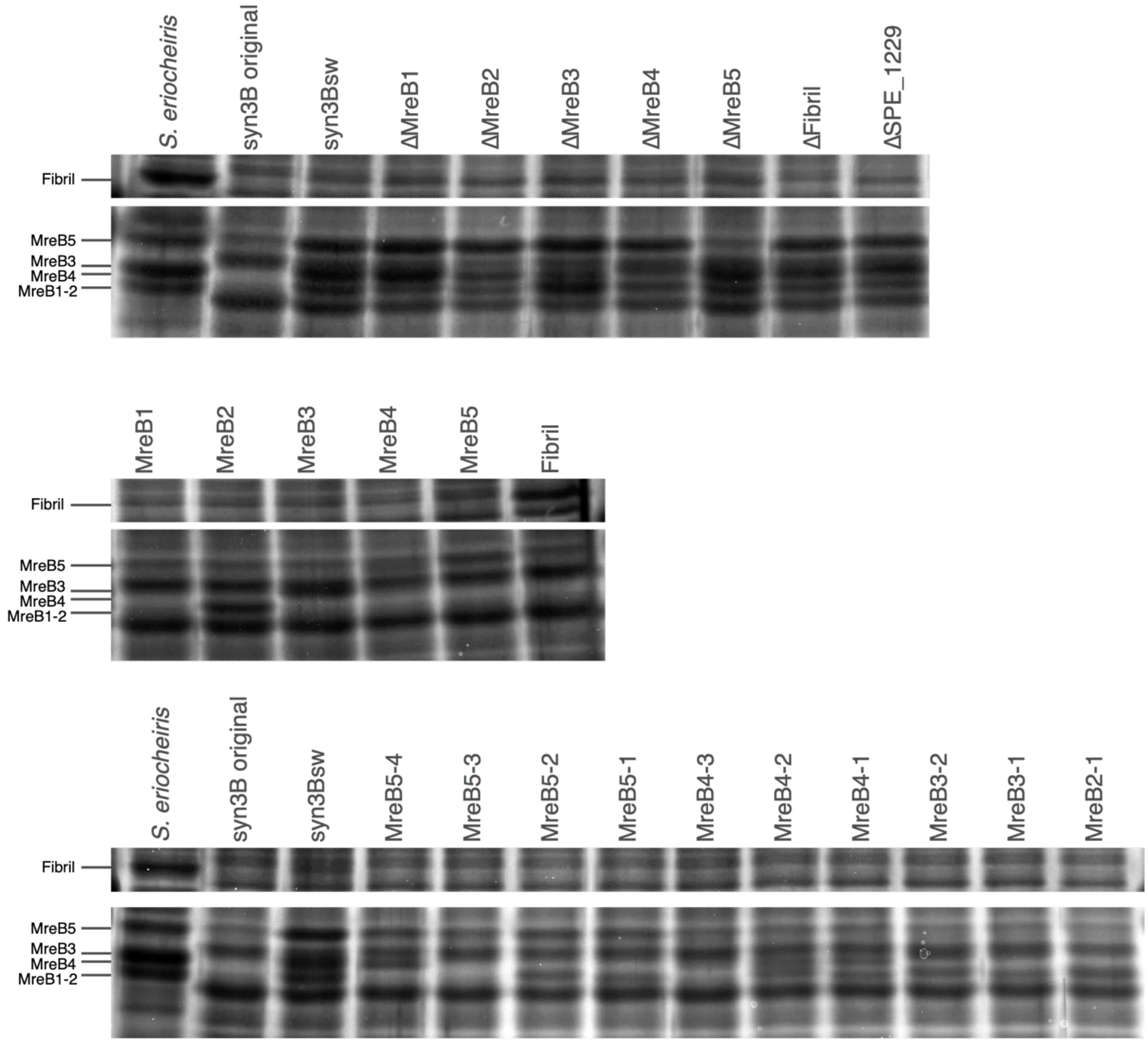


**Figure S5.**

Profiles of focused proteins visualized by SDS-10% PAGE. Constructs are indicated on the top, and the protein names are indicated on the left. Protein bands were stained by the reverse staining method.

**Table S1.**

DNA primers used in this study.

| Name | Sequence | Purpose |
|---|---|---|
| w_mreb3to5_R | TATAAAATTTAAAAATTACTTGGTATTGATAGTAAG | pSeW001 |
| w_mreb2_F | ACCAAGTAATTTTTAAATTTTATATTCTTTTGCTTC | pSeW001 |
| w_mreb2_R | TTAGATTATTAAATTCTCCCTCACTAAAACTAGTTG | pSeW001 |
| w_mreb1_F | GTGAGGGAGAATTTAATAATCTAATTCTTTGTTTTT | pSeW001 |
| w_mreb1_R | AAAAGTGAATAATTGCCTGAAGATTTAATAATAATT | pSeW001 |
| w_fibril_F | ATCTTCAGGCAATTATTCACTTTTTAAACGAATAGT | pSeW001 |
| w_fibril_R | TTGACAGCTAGCTCAGTCCTACCTTAAGTTTACTGGATTTTTAAG | pSeW001 |
| LP_inverse_R | GAACTATATAAATAACTCGCATATTG | pSeW001, pSemB1, pSemB2, pSemB3, pSemB4, pSemB5, pSefibril, pSeW435, pSeW425, pSeW415, pSeW325, pSeW315, pSeW215 |
| LP_inverse_F | AGGACTGAGCTAGCTGTCAAAGATC | pSeW001, pSemB2, pSefibril |
| w_mreb3to5_F | GCGAGTTATTTATATAGTTCTTATTTTTTCTTCCCAATGCCAGC | pSeW001, pSemB5 |
| addFib-to079-F | GAGTTATTTATATAGTTCTTATTCACTTTTTAAACGAATAGT | pSeW002, pSefibril |
| addFib-toPtuf-R | TTTTAAGGAGAAAAAAACATGATTGGAGTTATTTCAACTGCG | pSeW002, pSefibril |
| Ptuf-to079-R | GACAGCTAGCTCAGTCCTTATTTTTTGAATTAAGTATTAAAT | pSeW002, pSefibril |
| Ptuf-F | GTTTTTTTCTCCTTAAAATTTCTATAAC | pSeW002, pSefibril, pSeW545, pSeW535, pSeW525, pSeW515, pSeW435, pSeW425, pSeW415, pSeW325, pSeW315, pSeW215 |
| Stop-mreB1-F | TACAAAAGTCGGTTACTTATTGTTAATC | pSeW102 |
| Stop-mreB1-R | CAATAAGTAACCGACTTTTGTATCAATT | pSeW102 |
| Stop-mreB2-F | TACGCGACAGTTTATGCAGTTCCTAA | pSeW202 |
| Stop-mreB2-R | GAACTGCATAAACTGTCGCGTATGTC | pSeW202 |
| Stop-mreB3-F | ACGAGGTGGTTAAGGAGAAATATTAAATG | pSeW302 |
| Stop-mreB3-R | TTTCTCCTTAACCACCTCGTAAATTTATTG | pSeW302 |
| stop-mreB4-F | GAAAGTTGGTTATTTACTTTTGCCGCTA | pSeW402 |
| stop-mreB4-R | AAAGTAAATAACCAACTTTCGTTTCAAT | pSeW402 |
| Stop-mreB5-F2 | CCCTTGACCTTAAACGTAAGCTAACACG | pSeW502 |
| Stop-mreB5-R2 | CTTACGTTTAAGGTCAAGGGATTATCTA | pSeW502 |
| MreB1-F | GCGAGTTATTTATATAGTTCTTAATAATCTAATTCTTTGTTTTT | pSemB1 |
| MreB1-Pspi-R | GAGAAAGGAAATATAAGATCATGGCATTGATTAACAATAAGAAA | pSemB1 |
| Pspi_inverse-F | GATCTTATATTTCCTTTCTCTATT | pSemB1, pSemB3, pSemB4, pSemB5 |
| MreB2_R | ATGGCTAATTATAAATTTGGAAAA | pSemB2 |
| MreB2_Pspi_F | AAATTTATAATTAGCCATGATCTTATATTTCCTTTCTCTATTAAGTAG | pSemB2 |
| LP_insert_Pspi_R | GACAGCTAGCTCAGTCCTAATTAAAAGTTAGTGAACAAGAAA | pSemB2 |
| LP_insert_MreB2_F | GAGTTATTTATATAGTTCTTAAATTTTATATTCTTTTGCTTC | pSemB2, pSeW215 |
| Mreb3-Pspi-R | GAGAAAGGAAATATAAGATCATGACTATAACAGACGTATTAAAA | pSemB3 |
| MreB3-F | GCGAGTTATTTATATAGTTCTTATTTATTTTTTTTATTTTCTTC | pSemB3, pSeW325, pSeW315 |
| MreB4-Pspi-R | GAGAAAGGAAATATAAGATCATGTTAGATATTGTTTATGTTTAT | pSemB4 |
| MreB4-F | GCGAGTTATTTATATAGTTCTTAGTAATCTAATTCTTTAGTATG | pSemB4, pSeW435, pSeW425, pSeW415 |
| MreB5-Pspi-R | GAGAAAGGAAATATAAGATCGTGAAACCAGAAAGACCATTTATC | pSemB5 |
| mreB4-Ptuf-ORF-R | AATTTTAAGGAGAAAAAAACATGGCAGGATTTAATAGCGGCAA | pSeW545 |
| SD-mreB5-R | ATATTAAGGAGGAAATTAACGTGAA | pSeW535, pSeW525, pSeW515 |
| mreB3-Ptuf-R | AATTTTAAGGAGAAAAAAACATGACTATAACAGACGTATTAAAA | pSeW535, pSeW435 |
| mreB3-SD-mreB5-F | GTTAATTTCCTCCTTAATATTTATTTATTTTTTTTATTTTCTTCAATT | pSeW535 |
| mreB2- Ptuf-R | AATTTTAAGGAGAAAAAAACATGGCTAATTATAAATTTGGAAAA | pSeW525, pSeW425, pSeW325 |

| | | |
|---|---|---|
| mreB2- SD-mreB5-F | GTTAATTTCCTCCTTAATATTTAAATTTTATATTCTTTTGCTTC | pSeW525 |
| mreB1- Ptuf-R | AATTTTAAGGAGAAAAAAACATGGCATTGATTAACAATAAGAAA | pSeW515, pSeW415, pSeW315, pSeW215 |
| mreB1- SD-mreB5-F | GTTAATTTCCTCCTTAATATTTAATAATCTAATTCTTTGTTTTTG | pSeW515 |
| mreB3-mreB4-F | CCTCCCTTTTTTATTTATTTTTTTTATTTTCTTC | pSeW435 |
| mreB4-mreB3-R | AAATAAATAAAAAAGGGAGGAATTTTTACAATG | pSeW435 |
| mreB2_mreB4_F | CCTCCCTTTTTTAAATTTTATATTCTTTTGCTTC | pSeW425 |
| mreB2_mreB4_R | TAAAATTTAAAAAAGGGAGGAATTTTTACAAT | pSeW425 |
| mreB1-only-F | TTAATAATCTAATTCTTTGTTTTTGA | pSeW415, pSeW315 |
| mreB4-mreB1-R | ACAAAGAATTAGATTATTAAAAAAGGGAGGAATTTTTACAATG | pSeW415 |
| mreB3-mreB1-R | ACAAAGAATTAGATTATTAAATGAAAGGAGACAATCATAGATG | pSeW315 |
| mreB4-mChe-F | ATAATAGCTGATGATCCTGAGTATTTTGCTAATGAACCAATTTC | pSeW545-F4 |
| mreB4-mChe-R | ATAAAAGTGGAGCTCCTGGTCCAGACGAAAGAAAAATGAAAGTT | pSeW545-F4 |
| mreB5-mChe-F | ATAATAGCTGATGATCCTGAGTATTTAACTAATGAACCGATGTA | pSeW545-F5 |
| mreB5-mChe-R | ATAAAAGTGGAGCTCCTGGTCATAATGAACGTGCAATGCAAATT | pSeW545-F5 |
| SWmCh-linker-R2 | TCAGGATCATCAGCTATTATTAAAGAATTT | pSeW545-F5, pSeW545-F4 |
| SWmCh-linker-F | ACCAGGAGCTCCACTTTTATATAGTTCATCCATACCAC | pSeW545-F5, pSeW545-F4 |
| mreB4-SD-fibril-R | TTCGTTTAAAAAGTGAATAAAAAAGGGAGGAATTTTTACAATGG | pSeW6545 |
| fibril-F | TTATTCACTTTTTAAACGAATAGTTAC | pSeW6545 |
| coloP-puroR-F-Hana | ggagtagtccaaacagcaacagca | colony PCR |
| syn3B-junc-F | TATGTGATAATGCCAATCGCTAAG | colonyPCR |
| syn3B-junc-R | GTAAATTCCCAAATTATTCCATCA | colonyPCR |

**Table S2.**
Protein identification by mass spectrometry for Fig. S4.

| Bacterial strain | Protein band | Gene ID | Annotation | Mass (kDa) | score | coverage (%) |
|---|---|---|---|---|---|---|
| *S. eriocheiris* | i | SPE_0666 | Fibril | 58.7 | 184 | 63 |
| *S. eriocheiris* | ii | SPE_1231 | MreB5 | 38.7 | 117 | 54 |
| *S. eriocheiris* | iv | SPE_1230 | MreB4 | 40.7 | 64 | 49 |
| *S. eriocheiris* | iv | SPE_1049 | GAPDH | 35.8 | 129 | 68 |
| *S. eriocheiris* | v | SPE_1224 | MreB2 | 37.9 | 55 | 36 |
| original syn3B | i | ODP2_MYCCT | 2-oxo acid dehydrogenase subunit E2 | 47 | 43 | 24 |
| original syn3B | iii | - | - | - | - | - |
| syn3Bsw | i | SPE_0666 | Fibril | 58.7 | 124 | 52 |
| syn3Bsw | i | ODP2_MYCCT | 2-oxo acid dehydrogenase subunit E2 | 47 | 69 | 36 |
| syn3Bsw | ii | SPE_1231 | MreB5 | 38.7 | 131 | 57 |
| syn3Bsw | iii | SPE_1228 | MreB3 | 38.5 | 187 | 71 |
| syn3Bsw | iv | SPE_1230 | MreB4 | 40.7 | 86 | 57 |
| syn3Bsw | iv | SPE_1228 | MreB3 | 38.5 | 89 | 48 |
| syn3Bsw | iv | SPE_1224 | MreB2 | 37.9 | 46 | 34 |
| syn3Bsw | v | SPE_1224 | MreB2 | 37.9 | 101 | 42 |
| syn3Bsw | v | SPE_0470 | MreB1 | 38 | 96 | 45 |

**Movie S1.**
Cell behaviors of three strains indicated on top. Real time movie for 5 s.

**Movie S2.**
Rotational behaviors of freely moving site of *Spiroplasma* and syn3Bsw cells for 10 s.

**Movie S3.**
Cells lacking one of seven proteins from syn3sw for 10 s.

**Movie S4.**
Cells expressing a single *Spiroplasma* protein for 5 s.

**Movie S5.**
Syn3B cells expressing a pair of SMreBs for 10 s.

**Movie S6.**
Cells expressing a single protein for 10 s.

**Movie S7.**
SMreB5 localization in cell expressing SMreB 4 and 5 visualized by fluorescence for 10 s.

**Movie S8.**
syn3B cells expressing SMreBs 4, 5, and fibril for 10 s.

**Data S1. (separate file)**
DNA sequences of constructs used in this study.
